# Effector-dependent modulation of working memory maintenance in posterior parietal cortex

**DOI:** 10.1101/650077

**Authors:** Artur Pilacinski, Melanie S. Höller-Wallscheid, Axel Lindner

## Abstract

Working memory (WM) is the key process linking perception to action. Several lines of research have, accordingly, highlighted WM’s engagement in sensori-motor associations between retrospective stimuli and future behavior. Using human fMRI we investigated whether prior information about the effector used to report in a WM task would have an impact on the way the same sensory stimulus is maintained in memory – even if a behavioral response could not be readily planned. Specifically, we focused on WM-related activity in posterior parietal cortex during the maintenance of spatial items for a subsequent match-to-sample comparison, which was reported either with a verbal or with a manual response. We expected WM activity to be higher for manual response trials, because of posterior parietal cortex’s engagement in both spatial WM and hand movement preparation. Increased fMRI activity for manual response trials in bilateral anterior intraparietal sulcus confirmed our expectations. These results imply that the maintenance of sensory material in WM is optimized for motor context of the upcoming behavioral responses.

## INTRODUCTION

Working memory (WM) is the key cognitive process that allows bridging between previously encountered sensory information and future action. Yet, the detailed WM processing architecture and its underlying neuronal substrates still remain somewhat elusive [1, 2]. The traditional view on working memory architecture, which builds on Baddeley’s and Hitch’s multicomponent model, thereby assumes that the sensory information is maintained within domain-specific modules, namely the visuomotor sketchpad, the phonological loop, or the episodic buffer [3]. Brain imaging studies partially support this assumption, showing that WM-processing of a specific content engages the sites of its respective cortical representation, such as fusiform face area for faces, visual cortex for other visual forms, and temporal cortex for auditory content, [4, 5, 6, 7]. These content-specific WM “storages” possess independent processing capacities, further supporting the notion of separate storage modules for stimulus maintenance [8, 9].

While the classical concept of separate memory storage modules relates to the retrospective aspect of WM, i.e. it focuses on the content of the information that has been received, one should keep in mind that the information is maintained within WM in order to be used for future action. These latter, prospective aspects of WM and their link to motor behavior have been at the center of various investigations [10, 11, 12, 13, 14]. For example, rather than merely storing retrospective stimulus material, it has been shown that subjects do form and maintain prospective plans for actions in WM whenever possible [11, 14, 15].

Figure 1 A) and B) schematically illustrate typical experimental settings which highlight the maintenance of retrospective vs. retrospective information in WM. Figure 1 A) represents a “classical” working memory task (such as the Sternberg task [16]), which usually does not allow for action preparation during WM maintenance. Accordingly, only retrospective stimulus content should be maintained in WM. Figure 1 B) represents a task in which the task rule additionally allows a subject to readily plan and maintain an action. Accordingly, it might be the prospective action plan that is (additionally) maintained in WM in such situation. Experimental support has been provided in support of this conceptual distinction between retrospective and prospective information maintenance: Studies of Curtis and colleagues [11, 17] and Lindner and coworkers [14] report differences in brain activity whenever spatial objects not only had to be remembered but were, at the same time, targets for an upcoming goal directed actions such as saccades [11] or reaches [14]. Yet, in spite of these reported differences, both working memory and action planning tasks recruit an almost identical set of posterior parietal and frontal areas, reflecting strong functional ties between the two aspects of the perception-to-action processing stream [2, 11, 14, 18].

**Figure 1.**
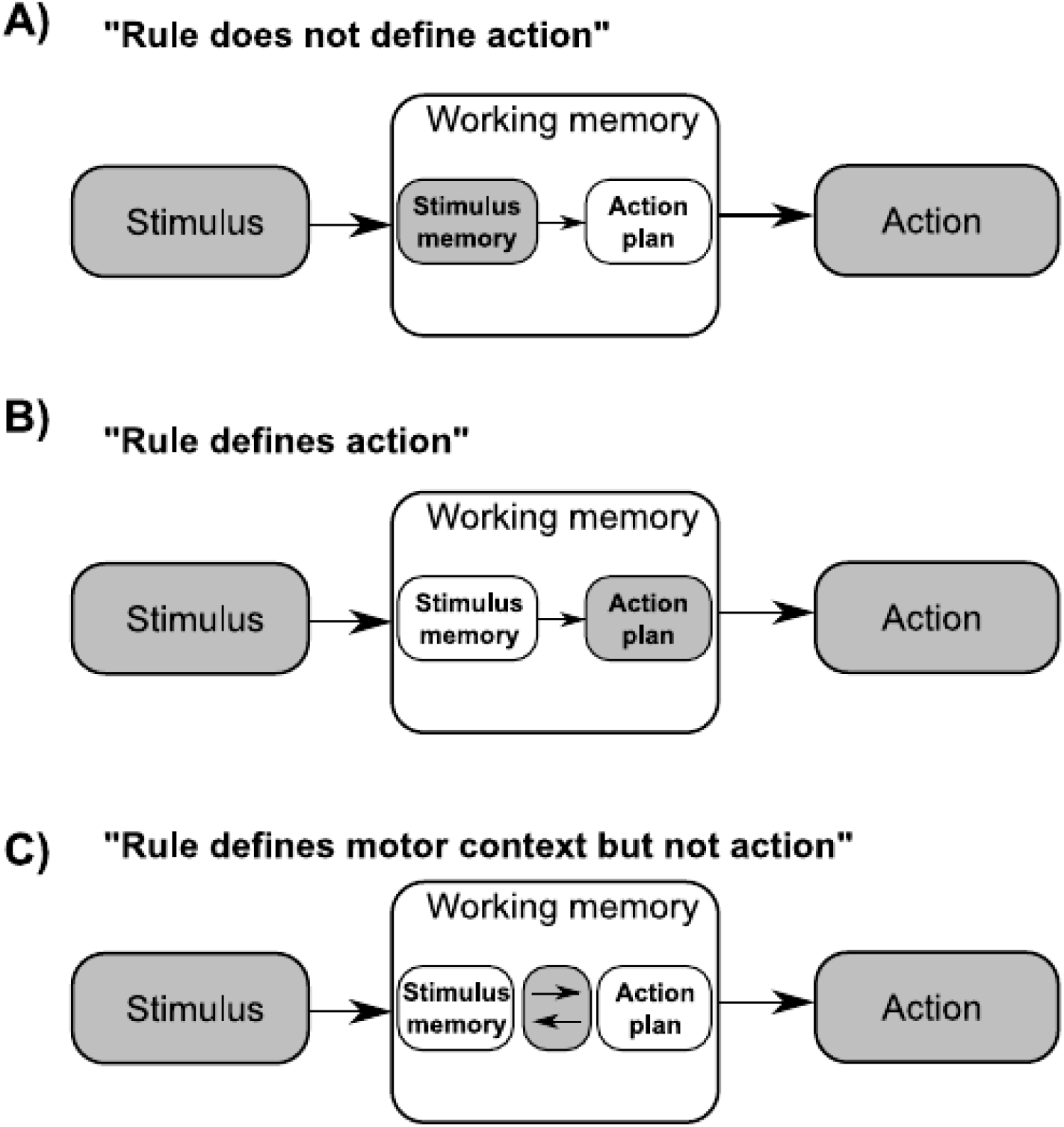
Hypothetical influence of varying task rules (A-C) on the maintenance of information in working memory. For further explanation, please refer to the main text.

This tangled relation between prospective and retrospective WM maintenance might be surprising at first glance. Yet, in natural environments the retrospective memory content typically predicts the motor context in which the memorized information will be used as both are naturally linked. For example, in natural world the probability that visuospatial information informs oculomotor or skeletomotor actions (e.g. saccades or reaches, respectively) is usually much higher than its probability for informing a verbal report. It follows that retrospective WM content could be maintained in a way that is optimized for the type of action that the memorized information affords (e.g. eye and hand movements in case of visuospatial information, etc.) – even in situations in which the ultimate action cannot be readily planned. A somewhat different rationale holds true for experimental settings in that even those WM tasks, which exclusively probe for the maintenance of retrospective sensory information (and which prohibit action planning), do typically require a predefined effector (such as pressing a button with one’s hand, etc.). Can the effector, which is afforded by a sensory stimulus or by a task rule, already have an impact on the way retrospective sensory information is maintained in working memory?

To answer this question, we devised a WM experiment where subjects’ brain activity was measured by means of functional magnetic resonance imaging (fMRI). Our experimental task was designed in a way that allowed us to eliminate processes related to specific action planning but to assess effector-dependent changes in brain activity representing WM maintenance of the same visuospatial stimuli. We expected that the sustained BOLD signals representing working memory maintenance may be influenced by effector modality through processes interrelating the retrospective stimulus memory and prospective motor actions (Figure 1C). These processes include the specification of the effector, which is afforded by the stimulus/task, as the main distinctive feature of the upcoming motor program [19]. Effector specification would, in turn, enable an optimization of memory storage to account for the expected/instructed future use of the memorized sensory material.

More specifically, we expected that the maintenance of visuospatial information in WM should lead to increased levels of fMRI activity in posterior parietal cortex (PPC) whenever the WM information will be probed with a hand response as compared to a verbal response. This is for a number of reasons. First, PPC has been demonstrated to be engaged both in visuospatial working memory tasks as well as in visuospatial planning tasks that engage eye and hand movements [11, 14, 20]. Second, PPC is less engaged in verbal preparation than frontal speech areas [21] and, moreover, it contributes significantly less to verbal as compared to spatial memory tasks [6]. Therefore, if WM maintenance of visuospatial information would be influenced by motor context (i.e. the instructed effector), we’d expect stronger activity in PPC in case of hand than in case of verbal responses.

In other words, if there is any influence of motor context on the maintenance of retrospective sensory information in WM, we should be able to trace it as an effector-dependent BOLD-signal modulation in PPC.

## RESULTS

Our experiment was comprised of two basic paradigms: a working memory, match-to-sample task (M2ST) and a control task (CT). The first one was a classical working memory paradigm with delayed response, where subjects were required to remember a visuospatial pattern of either two or otherwise six circles on the screen (presentation time: 3 sec). Then, after a 14/15 sec delay, a second pattern with the same number of circles came up and subjects had to indicate whether the pattern had changed or not (“same” vs. “different”). Crucially, in each trial we instructed the subjects immediately before the presentation of the circle stimuli to answer either manually (by using a button box held in their right hand) or verbally (compare: Figure 2; see “Materials and methods” for details). This instruction allowed us to manipulate the effector modality, which was relevant for the later behavioral response, on a trial by trial basis in both the M2ST and the CT. In the control task (CT) we also manipulated the instructed effector, but the subjects’ role was simply to judge the visual symmetry of circles presented after the delay (“symmetric” vs. “asymmetric”). Hence, the CT instructed the same effectors as in the M2ST but, importantly, did not require subjects to maintain spatial information in WM throughout the delay. The control task thus allowed us to account for any delay-related brain activity in the M2ST that would be merely related to anticipatory preparation of an effector as compared to a true reorganization of WM maintenance through the effector instruction. It is important to stress, again, that our experimental task ensured independence between the visuospatial memory content (i.e. the memory cue locations) and the specifically required motor responses (i.e. binary button presses).

**Figure 2.**
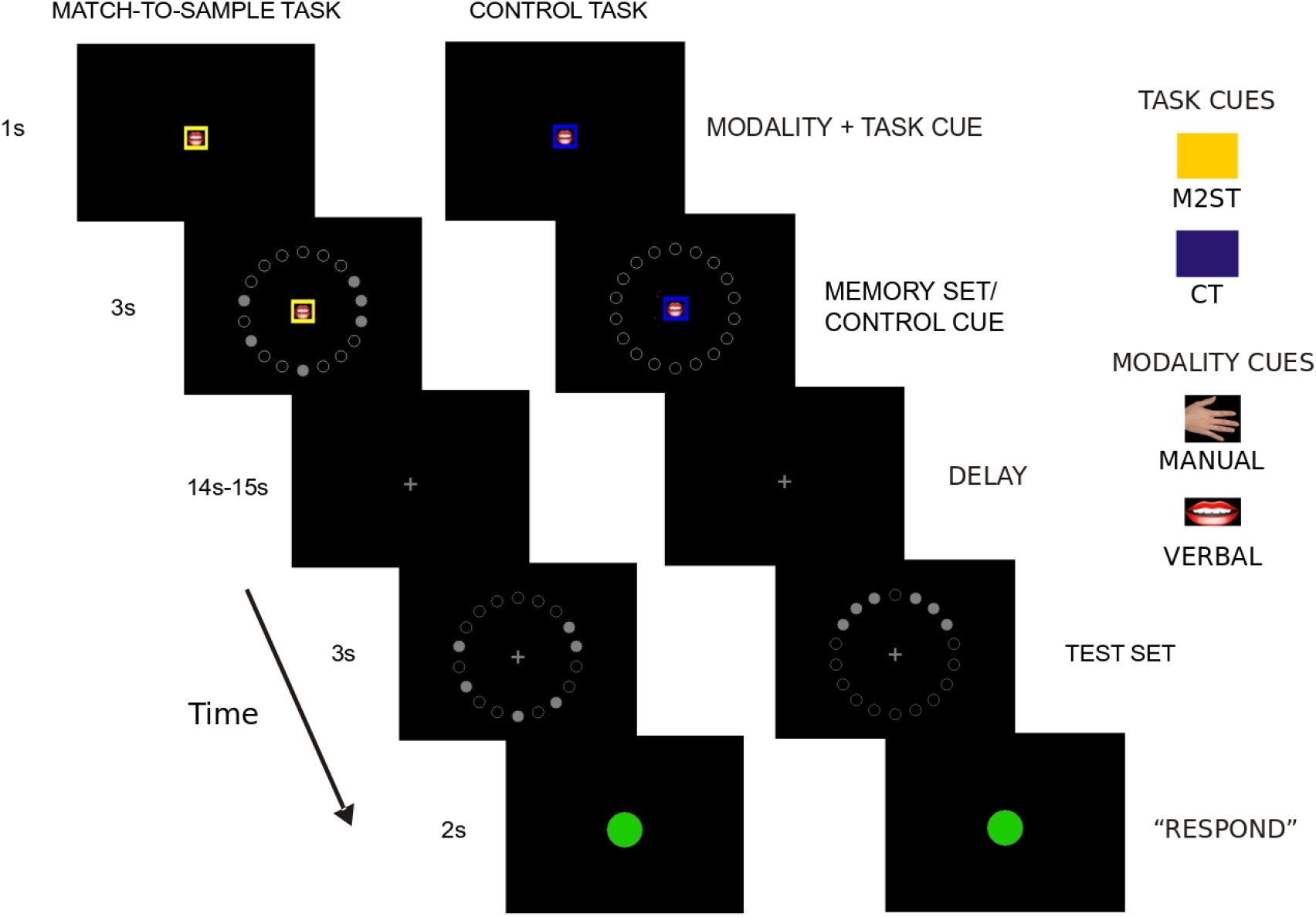
Exemplary timelines for our working memory task (M2ST) and the control task (CT). Each trial started with a color cue and an icon symbolizing the current task (M2ST or CT) and the response modality (Manual or Verbal), respectively. After displaying a memory item set (in M2ST) or, alternatively, an irrelevant control cue (in CT), the memory delay began. After the delay, a test set appeared, defining subjects motor response. In the M2ST subjects were asked to determine whether the presented test set matched the earlier memory set. In the CT subjects were asked to indicate whether or not the test set pattern was symmetric. A green “go” cue instructed the onset of the response period, in which subjects had to provide their answers in the correct response modality (manual or verbal). Please note that the visual mask screens were removed from the timeline for clarity. See text for further details.

Fourteen volunteers took part in the experiment. All of them were right handed, had no history of neurological disease and had normal or corrected to normal vision (see “Materials and methods” for details). All volunteers gave their written informed consent prior to the experiment, the experimental procedures were carried out in accordance with the declaration of Helsinki and the study was approved by the ethics committee at the University Clinic and the Medical Faculty of the University of Tübingen. Imaging was performed using a 3-Tesla MR-scanner and a twelve channel head coil (Siemens TRIO, Erlangen, Germany). Analyses were performed using SPM8 (Wellcome Center for Neuroimaging), R (R Foundation for Scientific Computing) and custom Matlab (MathWorks) routines.

We first analyzed the subjects’ task performance using a 2×2×2 ANOVA with the factors “task” (M2ST vs. CT), “response modality” (manual vs. verbal), and “load” (2 vs. 6 circles). Analyses of hit rates performed across all conditions revealed a significantly higher accuracy in the control task as compared to the memory task (see Figure 3; main effect of task: df=13, F=20.73, p=0.00054, eta^2^_G_=0.13). All other effects were not significant (load: df=13, F=0.47, p=0.5, eta^2^_G_=0.004; response modality: df=13, F=1.15, p=0.3, eta^2^_G_=0.008; modality x task: df=13, F=0.7, p=0.4, eta^2^_G_=0.005; response modality x load: df=13, F=02, p=.9, eta^2^_G_<0.001; task x load: df=13, F=2.99, p=0.01, eta^2^_G_=0.026; task x load x response modality: df=13, F=0.21, p=0.65, eta^2^_G_<0.001).

**Figure 3.**
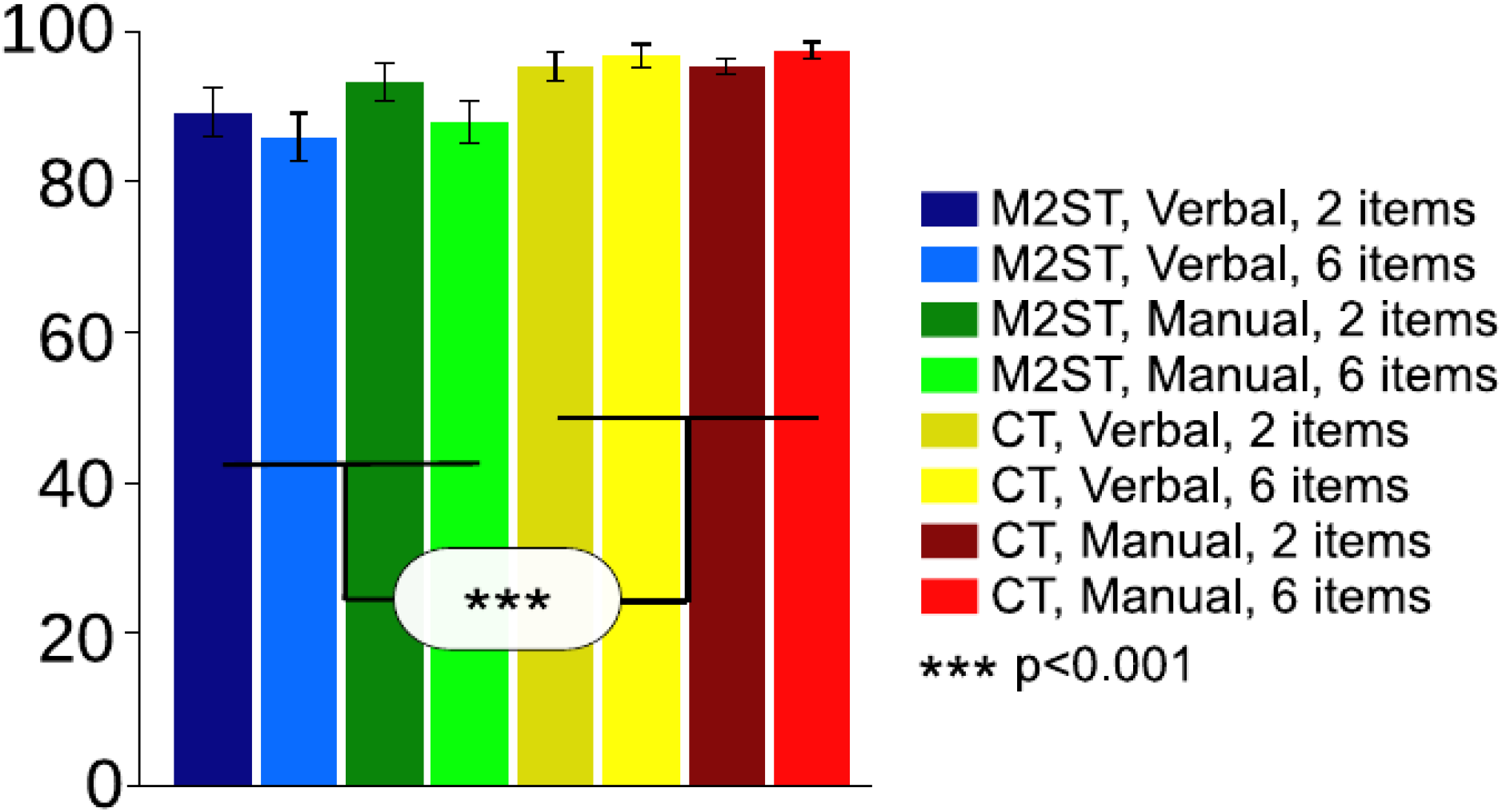
Task performance. The figure depicts across-subject average hit-rates (% correct) in the manual and the verbal response modality and separately for each load (2 or 6 items) and for each task (M2ST and CT). Error bars denote standard error. Hit rates differed significantly between M2ST and CT (for detailed statistics, please refer to the Results section).

Next, we quantified brain activity during the delay period of our tasks. To this end we used SPM 8, in which we specified a GLM for each subject. The GLMs comprised of eight conditions (2 tasks [M2ST & CT] x 2 response modalities [verbal & manual] x 2 ‘loads’ [2 and 6 circles]) that were modeled as separate boxcar regressors of respective duration for each of our trial epochs (cue presentation+mask, delay period, response phase; compare Figure 2). All regressors were convolved with the canonical haemodynamic response function in SPM8. Fixation epochs weren’t modeled explicitly and served as an implicit baseline. Note that we modeled the factor ‘load’ also in the control condition (as was defined by the number of symmetry test items visible in the response phase) despite this variable could not have any influence on brain activity in the preceding delay period. Accordingly, we did not consider load as a factor whenever analyzing delay-related brain activity in CT.

Figure 4 shows fMRI-activity increases during the delay phase in the M2ST as compared to the CT as revealed by an individual GLM-analysis in a representative subject (A) and by a second-level group-analysis (B). Such signal increases between tasks are expected due to the additional WM processes in M2ST as compared to CT. As expected, the M2ST yielded higher delay-related fMRI-activity in a number of fronto-parietal areas, which are typically recruited in working memory and action planning tasks [14]. These areas were bilateral superior parietal lobule (SPL), bilateral anterior intraparietal sulcus (aIPS), bilateral dorsal premotor cortex (PMd), bilateral ventral premotor cortex (PMv), bilateral dorsolateral prefrontal cortex (DLFPC) and supplementary motor area (SMA).

**Figure 4.**
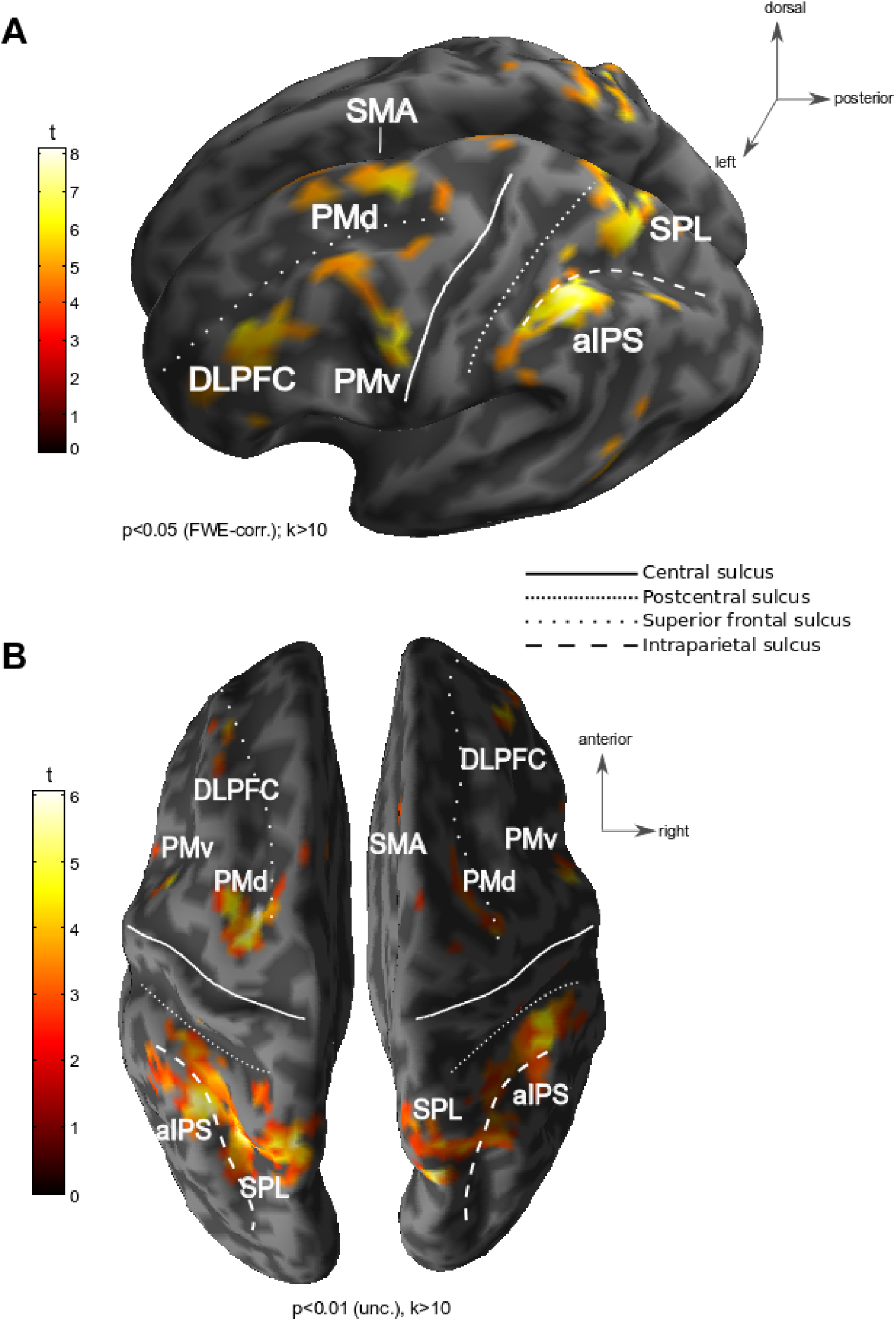
Working memory areas. The figures depict WM areas exhibiting stronger activity in M2ST than in CT during the delay period. A) depicts an exemplary subject’s WM contrast map. Such individual maps were used to functionally define our WM ROIs. B) depicts the WM contrast map of a corresponding random-effects group analysis.

All aforementioned areas were treated as regions-of-interest (ROIs) in our subsequent analyses. In these ROI analyses we probed for an influence of the instructed effector on delay-related activity in the M2ST as compared to the CT. Please note that we expected to reveal such influence in ROIs within posterior parietal cortex, as was laid out in detail in the introduction. Yet, to provide a more detailed overview, we also provide explanatory analyses of all other WM-ROIs. Moreover, we additionally included hand (M1h) and mouth (M1m) representations in the left primary motor cortex as control ROIs (see also “Materials and Methods” for further details and see Supplementary Table 1 for MNI coordinates of all ROIs).

From all aforementioned ROIs we extracted the beta weights of all delay-related GLM regressors in each individual. Beta values were expressed in terms of % signal change by dividing them through the overall residual beta estimates (i.e. our implicit baseline measure; compare above). The across-subject averages of these ‘normalized’ beta weights (collapsed across the two ‘loads’) are depicted in Figure 5. Visual inspection of the posterior parietal ROIs in this figure (upper row in Figure 5) reveals the expected activity pattern: First, there are higher levels of activity in the M2ST as compared to the CT (as was the defining criterion for ROI selection). To recapitulate, this was expected due to a contribution of PPC to visuospatial WM. Second, and more importantly, the delay-related activity was higher for instructed hand than for verbal responses in the M2ST but not in the CT.

**Figure 5.**
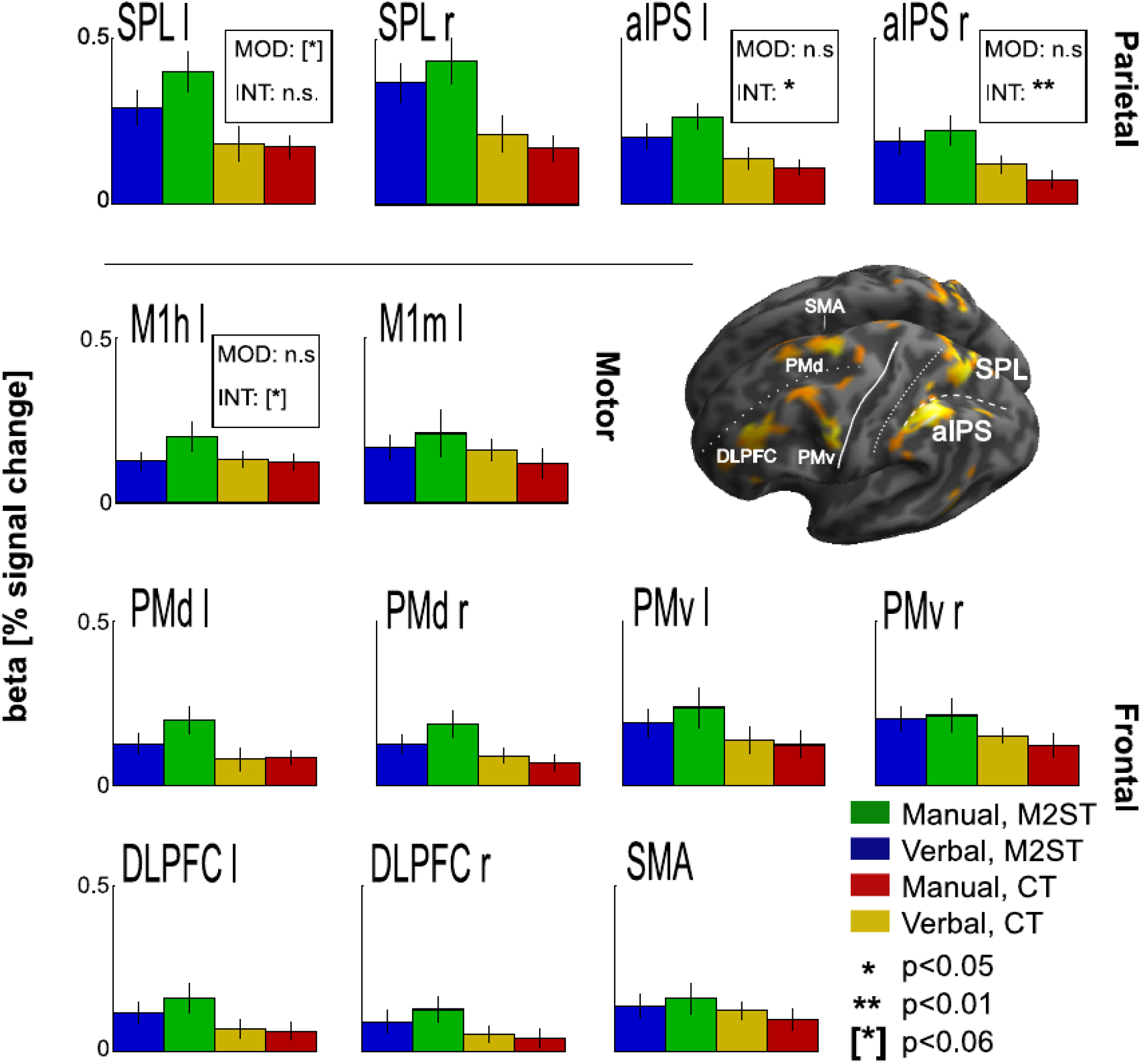
WM activity as a function of response modality and task. Individual bars reflect across-subject averages of delay-related beta estimates +/− SEM, averaged across both load conditions. Blue and green colors denote verbal and manual response modalities of M2ST, respectively. Yellow and red colors denote verbal and manual responses in CT. Additional statistical information from a corresponding 2×2 repeated measures ANOVAs with the factors “Response Modality” (MOD: manual vs. verbal) and “Task” (match-to-sample task vs. control task) is provided (for details, please refer to RESULTS). Significant signal differences between “verbal” and “manual” trials in M2ST as compared to CT (i.e. a significant interaction [INT]) were present in bilateral aIPS. This influence of response modality on maintenance-related brain activity was characterized by higher fMRI signal amplitudes in the M2ST whenever a hand response was prepared.

To statistically analyze these data, which seemingly matched our experimental hypothesis at first glance, we performed 2×2 ANOVAs (factors “Task” and “Response Modality”). This allowed us to directly compare delay-related activity in M2ST vs. CT as a function of effector modality and, thereby, to control for any unspecific motor readiness common to both tasks [22, 23]. If our hypothesis were true, we’d expect to reveal an interaction between “Task” and “Response Modality” at least in posterior parietal ROIs.

Indeed, our analyses confirmed such effector-dependent differences in fMRI-activity between M2ST and CT in the posterior parietal area aIPS bilaterally (“Response Modality”x”Task”. aIPS left: df=13, F=8.06, p=0.013, eta^2^_G_=0.03; aIPS right: df=13, F=10.14, p=0.0072, eta^2^_G_=0.02). Noteworthy, a trend for the same effect was also present in the left M1 hand area (df=13, F=4.3, p=0.059, eta^2^_G_=0.03).

Finally, we performed additional 2×2 ANOVAs to assess any potential influence of memory load on delay-related brain activity in the M2ST (factors “Response Modality” and “Load”). Please note that this analysis was restricted to the M2ST, because the factor “Load: had no impact on the respective activity estimates of CT: In the latter task, the factor load is merely relevant for the response period. The corresponding results of these additional analyses are provided as a supplement (compare supplementary table 2 and supplementary figure S1).

## DISCUSSION

In our study we observed that changes in working-memory-related BOLD activity in anterior IPS were influenced by the effector used for subjects’ subsequent response. This effect was not present in the control task, hence it is not merely attributable to “unspecific” effector preparation (or “motor readiness”). The major implication from these results is that the future motor context in which the memorized information is going to be reproduced, has an influence on memory maintenance. This principal finding offers a new perspective on the relationship between prospective and retrospective processes that constitute working memory.

It is important to emphasize, again, that our findings do not reflect a general, unspecific preparation of an effector or action [13, 24, 25], as was controlled for by our control task. We also ensured through our task design that there is a clear separation between the memory maintenance throughout the delay period and the preparation of any specific motor action, a problem that has been extensively highlighted in other works [10, 11, 17]: in our experiment, an action could only be planned and executed after the delay when the test set was shown. This means that the observed activity differences did not simply reflect effector-specific action planning during the delay phase. Instead, we show that sustained spatial working memory signals in the posterior parietal cortex were indeed affected by effector modality. Given that this activity was neither related to the planning of a specific action nor to unspecific action preparation, we postulate that it was the retrospective maintenance of the mnemonic material that is modulated by the future motor context of reproduction. In Figure 6 we suggest a hypothetical organization of such a feedback-based modulation of WM through the motor context as is afforded by the stimulus material or, as in our experiment, a task rule: These “affordances” could allow the WM system to infer the “motor context” in which sensory information is likely to be used and to guarantee optimal storage in “working memory” for guiding future action. It is important to stress that our framework is distinct from the notion that object affordances could provide additional semantic clues to support the maintenance of objects in working memory [26, 27] as, in related studies, object affordances were usually independent of any upcoming behavioral response. Moreover, we’d like to highlight that our framework clearly extends the multicomponent model of working memory, as was laid out in the introduction. Instead, it seems to be rather in agreement with more recent “state-based models of working memory”, which – according to D’Esposito and Postle [1] – “assume that the allocation of attention to internal representations—whether semantic LTM (e.g., letters, digits, words), sensory, or motoric—underlies the short-term retention of information in working memory” (page 3). In the following part we will discuss our results in the context of previous research on the contributions of posterior parietal cortex to the maintenance of retrospective vs. prospective information, we derive practical consequences from our work, we acknowledge potential limitations of our approach, and outline questions for future research.

**Figure 6.**
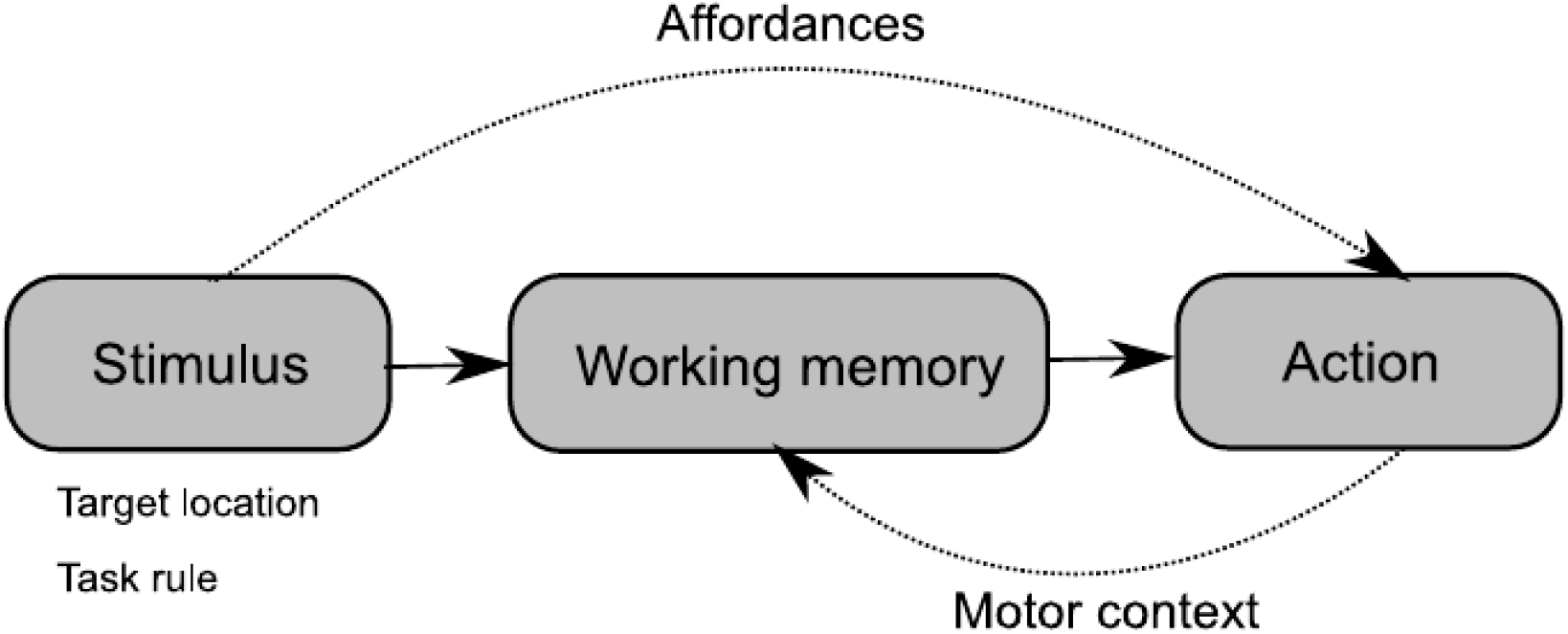
Hypothetical framework illustrating how the maintenance of mnemonic material could be modulated by future motor context. Both, mnemonic stimulus material (e.g. visual target location, as in our experiment) and task rules, could afford the motor context in which this material is likely to be used. In turn, motor context information could feed back on WM processes in order to guarantee optimal storage for guiding future action.

Our findings extend earlier research investigating the roles of posterior parietal and prefrontal cortex in prospective and retrospective working memory. For example, Curtis et al. disentangled the retro- and prospective processes in a visuospatial WM task engaging saccadic eye movements. They showed that the frontal oculomotor areas maintain prospective WM contents when subsequent eye movements should be directed at the remembered spatial location [11]. In turn, posterior parietal areas rather processed retrospective information when the remembered location was not a target for a subsequent saccade. On the downside, these authors could not exclude that their retrospective memory task – a non-match to sample task – required subjects to prospectively inhibit saccades to all remembered ‘non-target’ locations (which would be a type of prospective planning by itself). Following this argument, Lindner and colleagues [14], scrutinized the prospective (preparatory) role of PPC in planning: in a task that required both prospective and retrospective processing, PPC activity appeared to be driven mainly by prospective processes for both planning and inhibiting finger movements to visuospatial targets but, to a lesser extent, by retrospective memory, too. Our experiment extends these earlier findings by demonstrating a stronger involvement of posterior parietal cortex in visuospatial working memory maintenance whenever it involves the generation of a manual (rather than a verbal) response – even though the specific motor response could be neither planned nor inhibited.

Our results also have important practical implications: task-related brain activity in various working memory studies, which putatively did probe for maintenance of retrospective sensory information, might – at least in PPC – have been critically affected by the effector used for responding. This does not only call for a critical reconsideration of this earlier work. It does also call for a careful task design in future studies – one that takes the various potential contributions of behavioral components to working memory serious.

A limitation of our own study is that, so far, we have shown effector-dependent differences in WM maintenance only for spatial items and only in PPC, a region crucially involved in processing spatial WM content [11, 14, 20, 29]. On the basis of our current data we cannot determine whether a corresponding pattern of parietal activity, like the one reported here in parietal cortex for spatial content and manual responses, could be observed also for different memory modalities, e.g. the verbal content and verbal responses. It seems conceivable to think that patterns of activity would then generally shift towards areas representing the response effector that is congruent to the memorized material. For example, left lateral frontal cortex is specifically engaged both in the processing of verbal memory material [6] as well in the planning of verbal responses [21]. It is therefore likely that in this part of the brain one might likewise reveal an activity increase during the maintenance of verbal memory material. To scrutinize the detailed links between working memory and response preparation in other content and effector modalities than the ones presented here, more research is necessary.

In addition it would be interesting to see whether there is a true behavioral benefit from maintaining sensory information in a way that considers the relevant effector. The finding that eye- and hand- movements do more strongly affect the WM maintenance of spatial rather than verbal material may at least suggest that (spatial) WM maintenance does depend on congruent sensorimotor associations which is consistent with our rationale [28]. Unfortunately, our paradigm used here was not designed to probe for detailed behavioral effects. To this end we would have to use the verbal and manual response modality not only in combination with visuospatial memory material, but also in combination with verbal material. Only then we would be able to control for unspecific performance differences (e.g. reaction times, capacity estimates, or hit rates) between both the two effectors and the two memory modalities.

Lastly, although we expected a higher activity for manual responses in parietal regions, visual inspection of our data suggests that the presence of such expected pattern could be more widespread, and span also across frontal ROIs. This would not necessarily be surprising, as these ROIs have been previously attributed to hand action planning or to the maintenance of visuospatial memory too [6, 14]. Additionally intriguing is the apparent presence of effector-related modulation of activity in the hand area of primary motor cortex. Although that region has not exhibited any sustained delay-related response in the WM task, we can suspect that the effector-related processing in WM, as present in our task, did require some sort of information exchange with primary motor areas. At this very point we can only speculate about the actual nature of this exchange.

### Conclusions

In summary, our results suggest that the BOLD activity underlying working memory tasks is affected by the future motor context, namely the instructed effector. This effect was prominent in posterior parietal cortex, repeatedly demonstrated to be involved in both the maintenance of visuospatial information and the prospective planning of actions. Whether the posterior parietal cortex is indeed the key area bridging between the retrospectively processed mnemonic content and its future utilization, as is afforded by both the stimulus material and the task demands, remains to be answered. In any case, the top-down influence of a motor-preparatory component on the activity traditionally attributed to retrospective working memory maintenance sheds a new light on the actual structure of the working memory network and its intimate link to brain systems engaged in goal-directed actions.

## AUTHOR CONTRIBUTIONS

**A.P**.: designed the experiment, collected and analyzed data, wrote the manuscript. **M. H.-W**.: designed the experiment, collected fMRI and behavioral data, co-analyzed behavioral data, co-wrote “Materials and methods”. **A.L.**: co-wrote the manuscript, co-designed the experiment.

## MATERIALS AND METHODS

### Subjects

Fourteen subjects (two males; twelve females; mean age: 25) participated in this study. All of the participants had normal or corrected-to-normal visual acuity and were right-handed according to the Edinburgh Handedness Inventory [30]. None of them suffered from chronic, neurological or psychiatric diseases or took any medication. All subjects gave written informed consent prior to participation. Our experimental procedures were approved by the ethics committee at the University Clinic and the Medical Faculty of the University of Tübingen. Subjects received 10 Euro per hour for their participation. The group size was guided by power analyses performed on a similar, previously published fMRI dataset, comparing working memory and motor planning activity in posterior parietal cortex [14]. For a power of 0.80 and an alpha-level of 0.05 this analysis suggested a sample size of 11 subjects (two-tailed tests). To increase sensitivity, we initially decided to scan two further subjects. In the meantime, we have added yet another subject to our analysis in order to account for reviewer and editor requests.

### Task

We applied functional magnetic resonance imaging (fMRI) while our participants worked on either a delayed match-to-sample task in which subjects had to memorize dot patterns or a control task in which they had to judge if a given dot pattern is axially symmetrical (Fig. 2). Each trial started with a baseline period (14,000 or 15,000ms), in which subjects were asked to keep central fixation on a fixation cross. Then, a cue (1000ms) indicated a) if the current trial was a memory trial (yellow square) or a control trial (blue square) and b) if subjects would have to respond verbally (picture of a mouth) or manually (picture of a hand) in the end of the trial. In the memory trials, this cue period was followed by an encoding period (3000ms), in which subjects saw 18 small circles that were arranged in a circle around the central cue. Either two or six of these circles were filled, and their relative positions within the big circular arrangement served as the spatial memory items that subjects had to remember. Our spatial memory task design deliberately refrained from a sequential presentation of these visuospatial items. This was to avoid item ordering, which could affect WM processes of encoding and recall and, importantly, respective activity modulations in parietal regions that we could not easily control for [31]. The encoding screen was subsequently masked by a scrambled dot-pattern for 500ms to prevent afterimages of the cue and encoding screen. Then the delay epoch began (14,000 or 15,000ms). Afterwards, we either presented the very same dot pattern of the encoding period or a different one (3000ms). Subjects had to wait until they saw a green go-cue and had to indicate within 3000ms if this dot pattern did or did not match the one of the encoding period by either saying ‘same’ or ‘different’, respectively in the verbal conditions, or by pressing the right or left button of a button box, respectively, in the manual conditions. The control condition differed from the memory condition after the cue period insofar, as a circular arrangement consisting of 18 merely unfilled (but no filled) circles was presented so that subjects would not have to maintain any memory content during the subsequent delay period (14,000ms or 15,000ms). Then, we presented a circle composed of 18 small circles after the delay period. Again, either two or six of these circles were filled. Subjects waited until the green “go” cue and indicated whether or not this dot pattern was axially symmetrical to the vertical midline of the big circle by either saying “same” or “different”, respectively, in the verbal conditions or by pressing the right or left button of a button box, respectively, in the manual conditions. Lastly, a blank screen was displayed for the remaining 500ms of the trial. Our subjects worked on five consecutive blocks in which each of the 4 main conditions, i.e. 2 modalities (verbal vs. manual) x 2 tasks (memory vs. control), was presented twice and in a pseudo-randomized order, resulting in a total of 20 trials per condition for our main comparison M2ST vs. CT. Half of these trials (N=10) comprised of 2 items, the other half (N=10) comprised of 6 items.

### Stimulus presentation

We created the visual stimuli on a Windows^™^ based PC using MATLAB R2007b (The MathWorks, Inc.) and Cogent Graphics developed by John Romaya at the LON at the Wellcome Department of Imaging Neuroscience. They were projected onto a translucent screen (size of the projected image: 28 deg x 37 deg visual angle; viewing distance: 92 cm) by means of a video projector (frame rate: 60 Hz; resolution: 1024 x 768 pixels). Our participants watched the projected stimuli on the translucent screen being placed behind them with the aid of a mirror that was mounted on the head coil.

We displayed the fixation cross of the baseline and delay phases in Arial font and a 2.44 degrees visual angle font size. The squared color cue was 2.44 deg x 2.44 deg. The modality cue images were 1.95deg x 1.76deg. The spatial cues (dots) were placed at 4.9 deg radius from the central fixation point and were approximately 1 degree in diameter, each.

### Data acquisition

*Eye tracking.* Our subjects were supposed to maintain central fixation during the whole trial (besides the response phases) to ensure that fMRI activity would not be influenced by eye movements. We recorded eye-movements with a MRI-compatible infrared eye-camera (SMI SensoMotoric Instruments) and the ViewPoint Eye Tracker system (Arrington Research Inc.; sampling rate: 50 Hz) and performed on-line visual inspection of the eye image to ensure that subjects adhered to the fixation instruction. The proper detection of fixational saccades during scanning was not possible due to camera noise.

*WM performance.* In the verbal conditions, we recorded our subjects’ answers with a MRI-compatible microphone (Optoacoustics Dual-Channel Microphone, Optoacoustics Ltd., Israel; sampling rate: 8 kHz). The manual responses were recorded with a button box with two buttons indicating either “same” (left button) or “different” (right button). All recordings were analyzed off-line using self-written scripts in MATLAB R2007b (The MathWorks, Inc.). *fMRI data acquisition.* We collected the MR images with a 3-Tesla MR-scanner and a twelve channel head coil (Siemens TRIO, Erlangen, Germany). A T1-weighted magnetization-prepared rapid-acquisition gradient echo (MP-RAGE) structural scan of the whole brain was assessed from each subject (number of slices: 176, slice thickness: 1mm, gap size: 0 mm, in-plane voxel size: 1 x 1 mm, TR: 2300 ms, TE: 2.92 ms, FOV: 256 x 256 mm, resolution: 256 x 256 voxels). Moreover, we acquired T2*-weighted gradient-echo planar imaging (EPI) scans (slice thickness: 3.2 mm, gap size: 0.8 mm, in-plane voxel size: 3 x 3 mm, TR: 2000 ms, TE: 30 ms, flip angle: 90°, FOV: 192 x 192 mm, resolution: 64 x 64 voxels, 32 axial slices). 330 EPIs were collected from each participant during five consecutive blocks of 11 minutes each. Cerebral cortex and most sub-cortical structures were completely covered by the EPI-volume but we did not record from the most posterior parts of the cerebellum in several of our subjects due to brain size.

### Behavioral performance analysis

We statistically analyzed our behavioral data using SPSS (IBM SPSS Statistics, version 22) and R (R Foundation for Statistical Computing). Furthermore, functional MRI data were analyzed using SPM8 (Wellcome Department of Cognitive Neurology, London, UK) and R (R Foundation for Statistical Computing).

To investigate if the performance level was influenced by the modality, the task and/ or the load level, we analyzed share of correct answers by means of a three-way repeated measures ANOVA with the factors ‘response modality’ (2 levels: verbal vs. manual), ‘task’ (2 levels: memory vs. control), and ‘load’ (2 levels: 2 vs. 6).

### fMRI data analysis

Pre-processing. The pre-processing of our functional images was done in SPM8 (Wellcome Department of Cognitive Neurology, London, UK). Separately for each subject, we realigned all functional images by using the first scan of the first session as a reference. Then, we spatially coregistered the T1 anatomical image to the mean image of the functional scans and normalized our subjects’ mean anatomical images to the SPM T1 template in MNI space (Montreal Neurological Institute). The resulting normalization parameters were also applied to all functional images for spatial normalization. Finally, all functional images were smoothed by using a Gaussian filter (6 x 6 x 8 mm^3^ full-width at half-maximum) and high-pass filtered (cutoff period: 100 ms).

### First-level analysis

In the subject-level fMRI analysis we first specified a GLM for the M2ST with eight conditions (2 tasks x 2 response modalities x 2 loads) and three trial epochs each (cue+mask, delay, response), which were modeled as separate boxcar regressors (of corresponding epoch length), which were convolved with the canonical haemodynamic response function in SPM. Fixation epochs weren’t modeled explicitly and served in the model as an implicit baseline. In the control condition load served as a regressor of no interest in the cue+mask and in the delay epoch (defined as the number of symmetry test items visible in the response phase). Accordingly, for these two trial epochs the factor load was not considered in any of our analyses that would include the CT. Head motion estimates that were assessed during realignment were included in the model as six independent regressors (three regressors for head rotation and translation, each).

### Group-level analysis

Group-level activity maps were plotted on the basis of a 2^nd^ level random-effects analysis from the first level statistical parametric maps of activity related to the delay regressors in the CT and the M2ST (M2ST>CT). We thresholded statistical parametric maps at p<0.001, uncorrected for multiple comparisons, to ensure that our exploratory analysis of the whole brain does not omit any substantial activity during the delay phase. The resulting parametric maps were then overlayed on the standard MNI T1 template image as provided by SPM8 in order to anatomically define the regions of activity. Note that we used these maps only for visualization and aiding in selecting group-based coordinates for subsequent ROI selection in some individual subjects.

### ROI analysis

We focused the analyses on comparing activity in cortical areas involved in our working memory task. For the analysis of signal amplitudes, we selected areas in each individual subject, namely those that showed a statistically significant increase in BOLD intensity during the delay phase of working memory trials (as compared to control trials). We applied a statistical significance threshold of p<0.05, corrected for family-wise error. The maps were then overlaid on each subject’s T1 images, thus allowing precise assessment of the anatomical location of WM-related activation. Whenever this threshold did not yield any clusters of activation in a given subject in this full-brain analysis, we instead applied a small-volume correction centering on the coordinates of planning and spatial memory regions as were described by our group analyses (see Table S1). Small-volume correction was performed for a radius of 20mm around the respective ROI coordinates (p<0.05, FWE-corrected within this volume sphere). In one of the subjects, where the small-volume correction did not yield any significant clusters of activity, we used a t contrast comparing delay-related M2ST-activity against the implicit baseline (p<0.05, FWE-corrected). As we did not differentiate between the “response modality” at this stage of functional ROI definition (i.e. the activity maps were calculated for all conditions, pooled across both response modalities), our approach ensured that our ROI selection was not biased in favor of our hypothesis [32].

The ROIs were defined by the location of their corresponding local maxima of t values within major spatial clusters of working memory related BOLD activity. In particular, these were: left and right superior parietal lobule (SPL), left and right dorsal premotor cortex (PMd), left and right dorsolateral prefrontal cortex (DLPFC), left and right ventral premotor cortex (PMv), and left and right anterior intraparietal sulcus (aIPS). To complete our ROI set and allow observing any task-unspecific effector preparation we included anatomically defined functional representations of hand and vocal organs in the left primary motor cortex (labeled as M1h and M1m, respectively) [33, 34, 35].

**Table 1.**
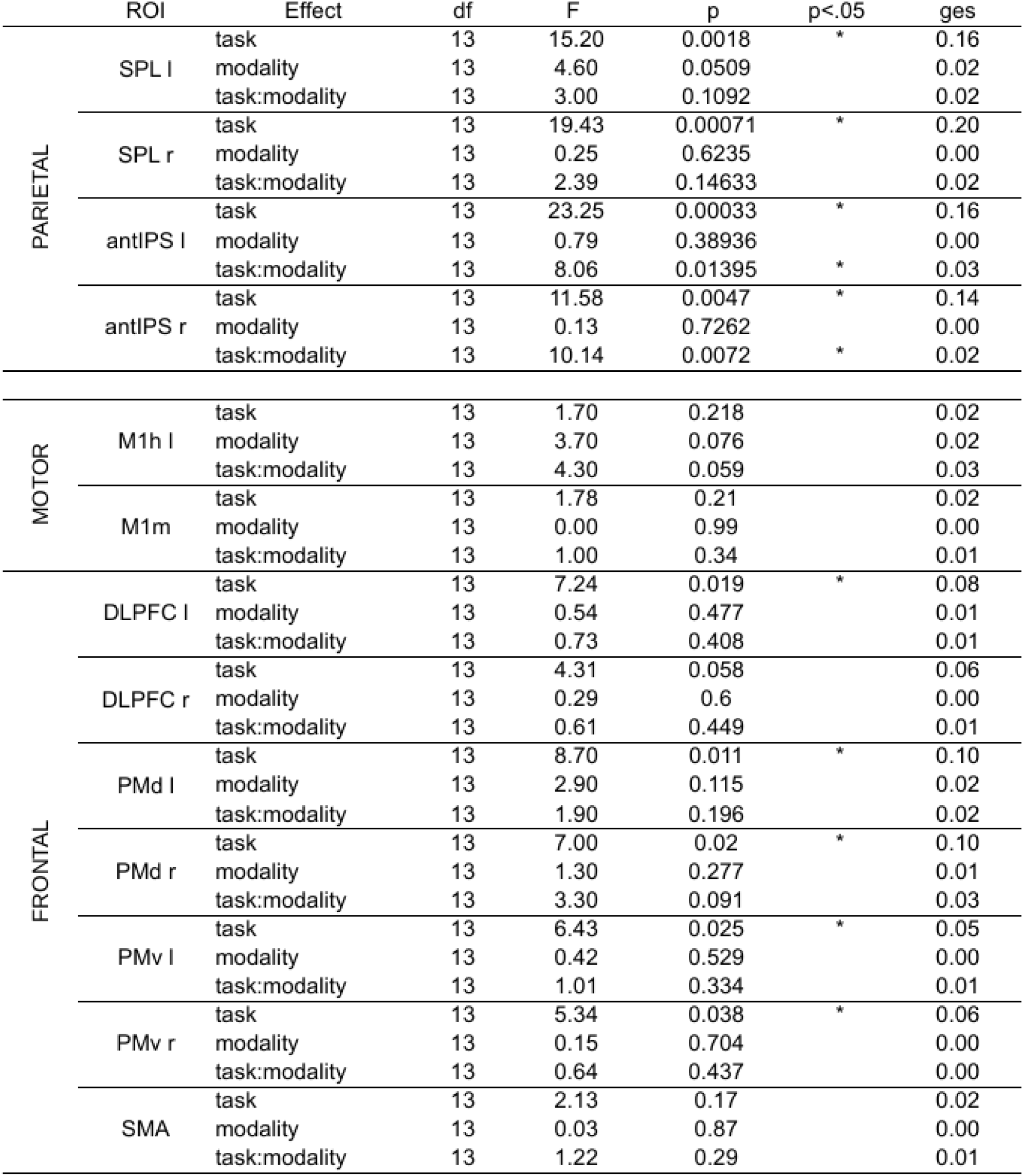
The table shows the results of our 2×2 repeated measures ANOVAs with the factors “Response Modality” (manual vs. verbal) and “Task” (match-to-sample task vs. control task), which was calculated across delay-related betas extracted from each ROI in each individual.

## REFERENCES

1. D’Esposito, M. & Postle, B. R. The Cognitive Neuroscience of Working Memory. Annu. Rev. Neurosci. 66, 115–142 (2015).

2. Eriksson, J., Vogel, E. K., Lansner, A., Bergström, F. & Nyberg, L. Neurocognitive Architecture of Working Memory. Neuron 88, 33–46 (2015).

3. Baddeley, A. The episodic buffer: a new component of working memory? Trends Cogn. Sci. 4, 417–423 (2000).

4. Ranganath, C., Cohen, M. X., Dam, C. & D’apos;Esposito, M. Inferior temporal, prefrontal, and hippocampal contributions to visual working memory maintenance and associative memory retrieval. J Neurosci 24, 3917–3925 (2004).

5. Harrison, S. A & Tong, F. Decoding reveals the contents of visual working memory in early visual areas. Nature 458, 632–5 (2009).

6. Wager, T. D. & Smith, E. E. Neuroimaging studies of working memory: A metaanalysis. Cogn. Affect. Behav. Neurosci. 3, 255–274 (2003).

7. Kirschen, M. P., Chen, S. H. A. & Desmond, J. E. Modality specific cerebro-cerebellar activations in verbal working memory: An fMRI study. Behav. Neurol. (2010).

8. Myerson, J., Hale, S., Rhee, S. H. & Jenkins, L. Selective Interference With Verbal and Spatial Working Memory in Young and Older Adults. Journals Gerontol. Ser. B Psychol. Sci. Soc. Sci. 54B, P161–P164 (1999).

9. Logie, R. H., Zucco, G. M. & Baddeley, A. D. Interference with visual short-term memory. Acta Psychol. (Amst). 75, 55–74 (1990).

10. Fuster, J. M. & Alexander, G. E. Neuron activity related to short-term memory. Science 173, 652–4 (1971).

11. Curtis, C. E. & D’Esposito, M. Selection and maintenance of saccade goals in the human frontal eye fields. J. Neurophysiol. 95, 3923–3927 (2006).

12. Leavitt, M. L., Mendoza-Halliday, D. & Martinez-Trujillo, J. C. Sustained Activity Encoding Working Memories: Not Fully Distributed. Trends Neurosci. 40, 328–346 (2017).

13. Curtis, C. E., Rao, V. Y. & Esposito, M. D. Maintenance of Spatial and Motor Codes during Oculomotor Delayed Response Tasks. 24, 3944–3952 (2004).

14. Lindner, A., Iyer, A., Kagan, I. & Andersen, R. a. Human Posterior Parietal Cortex Plans Where to Reach and What to Avoid. J. Neurosci. 30, 11715–11725 (2010).

15. van Ede, F., Chekroud, S. R., Stokes, M. G. & Nobre, A. C. Concurrent visual and motor selection during visual working memory guided action. Nat. Neurosci. 2, (2019).

16. Sternberg, S. Memory-scanning: mental processes revealed by reaction-time experiments. Am. Sci. 57, 421–57 (1969).

17. Mackey, W. E. & Curtis, C. E. Distinct contributions by frontal and parietal cortices support working memory. Sci. Rep. 7, 6188 (2017).

18. Gallivan, J. P. & Culham, J. C. ScienceDirect Neural coding within human brain areas involved in actions. Curr. Opin. Neurobiol. 33, 141–149 (2015).

19. Rosenbaum, D. A. Human movement initiation: specification of arm, direction, and extent. J. Exp. Psychol. Gen. 109, 444–474 (1980).

20. Pisella, L., Berberovic, N. & Mattingley, J. B. Impaired Working Memory for Location but not for Colour or Shape in Visual Neglect: a Comparison of Parietal and Non-Parietal Lesions. Cortex 40, 379–390 (2004).

21. Brendel, B. et al. The contribution of mesiofrontal cortex to the preparation and execution of repetitive syllable productions: An fMRI study. Neuroimage 50, 1219–1230 (2010).

22. Di Russo, F. et al. Beyond the “Bereitschaftspotential”: Action preparation behind cognitive functions. Neurosci. Biobehav. Rev. 78, 57–81 (2017).

23. Cui, X., Stetson, C., Montague, P. R. & Eagleman, D. M. Ready…go: Amplitude of the FMRI signal encodes expectation of cue arrival time. PLoS Biol. 7, e1000167 (2009).

24. Cavina-Pratesi, C. et al. Dissociating arbitrary stimulus-response mapping from movement planning during preparatory period: evidence from event-related functional magnetic resonance imaging. J. Neurosci. 26, 2704–13 (2006).

25. Connolly, J. D., Goodale, M. A., Menon, R. S. & Munoz, D. P. Human fMRI evidence for the neural correlates of preparatory set. Nat. Neurosci. 5, 1345–1352 (2002).

26. Mecklinger, A., Gruenewald, C., Weiskopf, N. & Doeller, C. F. Motor Affordance and its Role for Visual Working Memory: Evidence from fMRI studies. Exp. Psychol. 51, 258–269 (2004).

27. Pecher, D. No role for motor affordances in visual working memory. J. Exp. Psychol. Learn. Mem. Cogn. 39, 2–13 (2013).

28. Lawrence, B., Myerson, J., Oonk, H. & Abrams, R. The effects of eye and limb movements on working memory. Memory 9, 433–444 (2001).

29. Hamidi, M., Tononi, G. & Postle, B. R. Evaluating frontal and parietal contributions to spatial working memory with repetitive transcranial magnetic stimulation. Brain Res. 1230, 202–210 (2008).

30. Oldfield, R. C. The assessment and analysis of handedness: The Edinburgh inventory. Neuropsychologia 9, 97–113 (1971).

31. Majerus, S. et al. The left intraparietal sulcus and verbal short-term memory: focus of attention or serial order? Neuroimage 32, 880–91 (2006).

32. Kriegeskorte, N., Simmons, W. K., Bellgowan, P. S. F. & Baker, C. I. Circular analysis in systems neuroscience: the dangers of double dipping. Nat. Neurosci. 12, 535–40 (2009).

33. Yousry, T. A. et al. Localization of the motor hand area to a knob on the precentral gyrus. A new landmark. Brain 120, 141–157 (1997).

34. Simonyan, K. & Horwitz, B. Laryngeal motor cortex and control of speech in humans. Neuroscientist 17, 197–208 (2011).

35. Bouchard, K. E., Mesgarani, N., Johnson, K. & Chang, E. F. Functional organization of human sensorimotor cortex for speech articulation. Nature 495, 327–332 (2013).

